# Modeling of biomechanics and biorheology of red blood cells in type-2 diabetes mellitus

**DOI:** 10.1101/132134

**Authors:** Hung-Yu Chang, Xuejin Li, George Em Karniadakis

## Abstract

Erythrocytes in patients with type-2 diabetes mellitus (T2DM) are associated with reduced cell deformability and elevated blood viscosity, which contribute to impaired blood flow and other pathophysiological aspects of diabetes related vascular complications. In this study, by using a *two-component* red blood cell (RBC) model and systematic parameter variation, we perform detailed computational simulations to probe the alteration of the biomechanical, rheological and dynamic behavior of T2DM RBCs in response to morphological change and membrane stiffening. First, we examine the elastic response of T2DM RBCs subject to static tensile forcing and their viscoelastic relaxation response upon release of the stretching force. Second, we investigate the membrane fluctuations of T2DM RBCs and explore the effect of cell shape on the fluctuation amplitudes. Third, we subject the T2DM RBCs to shear flow and probe the effects of cell shape and effective membrane viscosity on their tank-treading movement. In addition, we model the cell dynamic behavior in a microfluidic channel with constriction and quantify the biorheological properties of individual T2DM RBCs. Finally, we simulate T2DM RBC suspensions under shear and compare the predicted viscosity with experimental measurements. Taken together these simulation results and their comparison with currently available experimental data are helpful in identifying a specific parametric model the first of its kind that best describes the main hallmarks of T2DM RBCs, which can be used in future simulation studies of hematologic complications of T2DM patients.

## INTRODUCTION

Diabetes mellitus (DM), the fastest growing chronic disease worldwide, is a metabolic dysfunction characterized by elevated blood glucose levels (hyperglycemia) (1). Type 2 DM (T2DM) is the most common form of diabetes. People with T2DM demonstrate significantly higher mortality rates relative to non-diabetics due to an increased risk of developing macro-and microvascular complications (2). Hemorheological abnormalities emerged in diabetic patients play a key role in the pathogenesis and progression of life-threatening coronary and peripheral artery diseases. One of the hemorheological determinants is the impaired deformability of red blood cells (RBCs) involved in T2DM (3). A healthy human RBC is a nucleus-free cell; it is primarily comprised of a fluid-like lipid bilayer contributing to the bending resistance, an attached spectrin network (cytoskeleton) maintaining cell shape and facilitating its motion, and transmembrane proteins bridging the connections between lipid and spectrin domains (4, 5). Owing to the fluid nature of the lipid bilayer and elastic nature of the cytoskeleton, the RBC is capable of dramatic deformations when passing through narrow capillaries as small as three microns in diameter without any damage.

For RBCs in pathological conditions, the alterations in cell geometry and membrane properties of diseased RBCs could lead to impaired functionality including loss of deformability (6–8). For example, the shape distortion and membrane stiffening of RBCs induced by parasitic infectious diseases like malaria (9, 10) and certain genetic blood disorders like sickle cell disease (11) cause increased cell rigidity and decreased cell deformability. The RBC deformability has also been demonstrated to be impaired in T2DM. For example, using micropipette aspiration and filtration techniques, McMillan *et al.* and Kowluru *et al.* showed that T2DM RBCs are less deformable and more fragile compared to non-diabetic subjects (12, 13). Agrawal *et al.* found that the size of T2DM RBC is larger than that of normal RBC because of the possible metabolic disturbances (14). Babu *et al.* showed that the development of irregularity in the contour of the T2DM RBCs under hyperglycemia would cause a significant reduction in cell deformability (15). In addition, several studies using atomic force microscopy (AFM) directly examined the biomechanical properties of diabetic RBCs and demonstrated that they are less deformable than normal RBCs (Fig. 1), which could be due to the oxidation and glycosylation of hemoglobin and proteins on cell membrane under abnormal glycemic condition (16–18).

**Figure 1:**
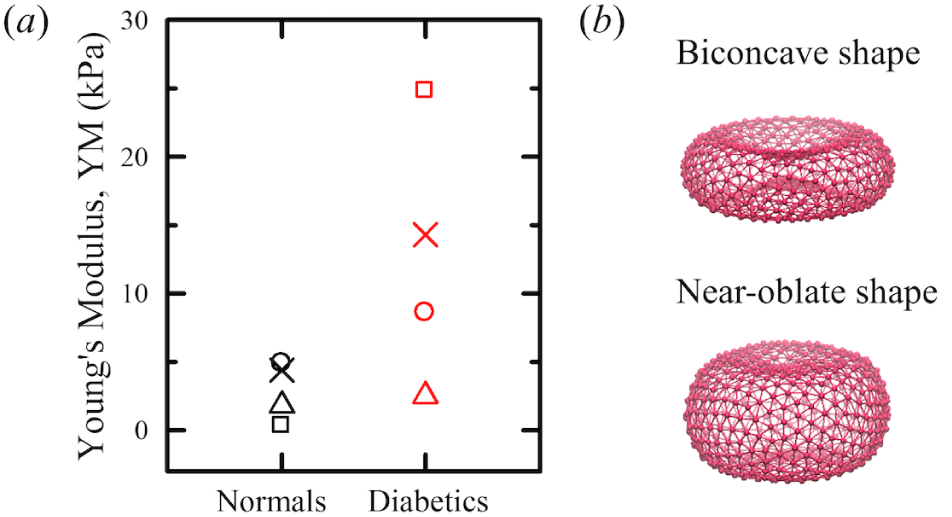
(a) Young’s modulus (YM) of normal and diabetic RBCs measured in experiments with data as follows: Cross (×), Fornal *et al.* (16); triangle (△), Ciasca *et al.* (17); square (◼), Zhang *et al.* (18); circle (◯), Lekka *et al.* (54). (b) Sketch of the RBC models with the equilibrium biconcave (*S/V* = 1.44) and near-oblate (*S/V* = 1.04) shapes.

The RBC deformability is an important hemorheological parameter in determining whole blood viscosity, hence blood flow resistance in the microcirculation (19–21). Increased blood viscosity has been demonstrated in patients with T2DM, which induces insufficient blood supply and vascular damage, and eventually leads to diabetic microangiopathy and other circulation problems. For example, Skovborg *et al.* revealed that the blood viscosity of the diabetic subjects is approximately 20% higher than that of controls (19). Ercan *et al.* suggested that elevated plasma cholesterol contributes to increased blood viscosity by an additional effect to hyperglycemia in T2DM patients (22). In long-standing DM with non-proliferative retinopathy, Turczynski *et al.* showed that the blood viscosity is positively correlated with the retinopathy severity (23), which is attributed to the decrease of RBC deformability.

Along with the aforementioned experimental studies, recent advances in computational modeling and simulation enable investigation of a broad range of biomechanical and rheological blood-related problems at different length scales (24–29). In recent years, mesoscopic particle-based RBC models, which treat both the fluid and the membrane as particulate materials, are increasingly popular as a promising tool for modeling the structural, mechanical, and rheological properties of RBCs in normal and pathological conditions (30**?**–33). Two different particle-based RBC models using dissipative particle dynamics (DPD) (34, 35), *i.e.*, the one-component RBC models (36–38) and the two-component RBC models (39), have been developed and employed to conduct efficient simulations of RBCs in the microcirculation (40–48). In particular, the two-component RBC model, which considers the lipid bilayer and cytoskeleton separately but also includes the transmembrane proteins, facilitates detailed whole-cell exploration of the diverse biophysical and biomechanical aspects involving RBCs, such as probing the multiple stiffening effects of nanoscale knobs on malaria-infected RBCs (10, 49), and predicting the biomechanical interactions between the lipid bilayer and the cytoskeleton in human RBCs (39, 50).

In the current study, we extend the two-component whole-cell model to T2DM RBCs and investigate the morphological and biomechanical characteristics of T2DM RBCs and hemorheological properties of T2DM blood. Specifically, we test and validate the whole-cell model through rigorous comparisons with experimental data from five different sets of independent measurements that probe different aspects of biomechanical and rheological properties of T2DM RBCs, including RBC stretching deformation and shape relaxation, membrane fluctuations, RBC dynamics in shear flow, and blood viscosity in T2DM. The rest of the paper includes section 2 for a brief description of the development of diabetic RBC models, section 3 for detailed simulation results and discussion, and section 4 for a summary of major findings and conclusion.

## METHODS

In this study, we probe the biomechanics, rheology and dynamics of T2DM RBCs with the help of the two-component RBC model based on the DPD method. The intracellular and extracellular fluids are modeled by collections of free DPD particles and their separation is enforced by bounce-back reflections at the RBC membrane surface. The RBC membrane interacts with the fluid and wall particles through DPD forces, and the system temperature is maintained by the DPD thermostat. For completeness, we review briefly the two-component RBC model below, whereas for detailed description of the DPD method and the RBC model, we refer to Refs. (34–37, 39).

### Two-component RBC model

In the two-component whole-cell model, the cell membrane is modeled by two distinct components, *i.e.*, the lipid bilayer and the cytoskeleton, and each component is constructed by a 2D triangulated network with *N*_*v*_ vertices. In general, the two-component RBC model takes into account the elastic energy, bending energy, bilayer-cytoskeleton interaction energy, and constraints of fixed surface area and enclosed volume, given by

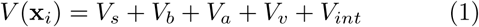

where *V*_*s*_ is the elastic energy that mimics the elastic spectrin network, given by

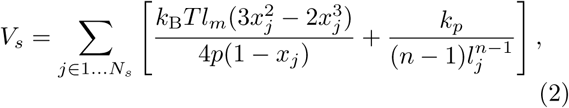

where *l*_*j*_ is the length of the spring *j*, *l*_*m*_ is the maximum spring extension, *x*_*j*_ = *l*_*j*_/*l*_*m*_, *p* is the persistence length, *k*_B_*T* is the energy unit, k_*p*_ is the spring constant, and *n* is a specified exponent. The bending resistance from the lipid bilayer of the RBC membrane is modeled by

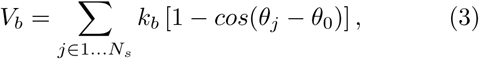

where *k*_*b*_ is the bending constant, *θ*_*j*_ is the instantaneous angle between two adjacent triangles having the common edge *j*, and *θ*_0_ is the spontaneous angle.

Constraints on the area and volume conservation of RBC are imposed to mimic the area-preserving lipid bilayer and the incompressible interior fluid. The corresponding energy is given by

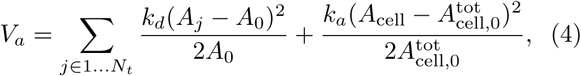

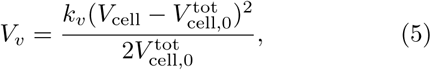

where *N*_*t*_ is the number of triangles in the mem brane network, *A*^0^ is the triangle area, and *k*_*d*_, *k*_*a*_ and *k*_*v*_ are the local area, global area and volume constraint coefficients, respectively. The terms 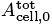 and 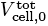 represent the specified total area and volume, respectively.

The bilayer-cytoskeleton interaction potential, *V*_*int*_, is expressed as a summation of harmonic potentials given by

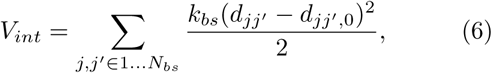

where *k*_*bs*_ and *N*_*bs*_ are the spring constant and the number of bond connections between the lipid bilayer and the cytoskeleton, respectively. *d*_*jj′*_ is the distance between the vertex *j* of the cytoskeleton and the corresponding projection point *j* ′ on the lipid bilayer, with the corresponding unit vector **n**_*jj′*_; *d*_*jj′, 0*_ is the initial distance between the vertex *j* and the point *j*′, which is set to zero in the current simulations.

### Mechanical properties of RBC membrane

It is known that the membrane elasticity of RBCs characterizes its resistance to deformation, and membrane viscosity characterizes the viscous resistance of the cell membrane to shear deformation (51). Following the linear analysis for a regular hexagonal network by Dao *et al.* (52), we correlate the model parameters and the network macroscopic elastic properties, *i.e.* shear modulus (*µ*), which is determined by

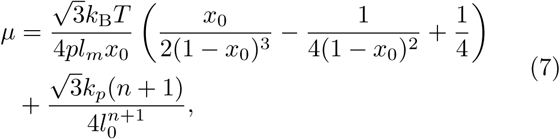

where *l*_0_ is the equilibrium spring length and *x*_0_ = *l*_0_*/l_m_*.

The relation between the modeled bending constant *k*_*b*_ and the macroscopic bending rigidity *k*_*c*_ of the Helfrich model can be derived as 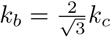 for a spherical membrane (53). This expression describes bending contribution of the energy in Eq. (3), but may not fully represent actual bending resistance of the RBC membrane as the membrane bending may also result in local in-plane deformations.

In addition, membrane viscosity also plays an important role in the dynamic behavior of RBC in physiological and pathological conditions. To estimate the effective membrane viscosity *η*_*m*_, we simply combine the viscous contributions from lipid bilayer *η*_*b*_ and cytoskeleton *η*_*s*_. These two viscous terms, following prior work (39),can be calculated as 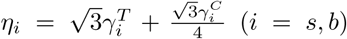, where 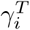 and 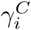 are dissipative parameters of the DPD model.

### Parameter estimation

Under physiological conditions, a normal RBC has a distinctive biconcave shape with a high surface area-to-volume (*S/V*) ratio, which facilitates the oxygen transport through the cell membrane and contributes to the remarkable cell deformability. In this study, we model a normal RBC (N-RBC) with the following parameters: total number of vertices *N*_*v*_ = 500, RBC membrane shear modulus *µ*_0_ = 4.73 *µ*N/m, bending modulus *k*_*c*_= 31.9 *k*_B_*T*, and effective membrane viscosity *η*_*m*_ = 0.128 *pa · s*, RBC surface area 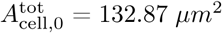, cell volume 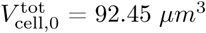, and surface area-to-volume ratio *S/V* = 1.44.

Different from the N-RBC, the diabetic RBCs have decreased cell deformability and increased cell volume. Here, we have developed three potential T2DM RBC models (D-RBC1, D-RBC2, and D-RBC3) based on previous *in vitro* experiments. First, we selected experimentally determined shear modulus data (55) and set the shear modulus of diabetic RBC model (D-RBC1) to *µ* = 2.0*µ*_0_. Second, previous AFM measurements suggest that there is a visible change in cell shape from the normal biconcave shape to a near-oblate shape with a reduced *S/V* ratio in the pathophysiology of T2DM RBCs (56). Considering this fact, here, we propose a modified diabetic RBC model (D-RBC2) with an oblate shape (*S/V ∼* 1.04) (Fig. 1b). Third, lower membrane fluidity and increased membrane viscosity for T2DM RBCs than controls have been reported and suggested to be the consequences of nonenzymatic glycation induced changes in the RBC membrane (57). In addition, previous studies also suggest a comparable shape recovery time (*t*_*c*_) between normal and diabetic RBCs (12, 58). It is known that the recovery time, *t*_*c*_ = *η*_*m*_/*µ*, is primarily determined by the viscoelastic properties of RBC membrane (51, 59). In combination with the above experimental results, we have developed a third diabetic RBC model (D-RBC3) by considering the aberrant cell shape and impaired deformability as well as adjusted effective membrane viscosity. In summary, the D-RBC1 model holds a biconcave shape (*S/V* = 1.44) as that of N-RBC with an increased *µ*, the DRBC2 model takes a near-oblate shape with a reduced *S/V* ratio (*∼* 1.04) accompanied by the increase in *µ*, and the D-RBC3 model has the same characteristics as that of D-RBC2 but the additional trait of enhanced *η*_*m*_. Parameters related to the membrane properties among the different RBC models, including the shear modulus *µ*, effective membrane viscosity *η*_*m*_, and surface area-to-volume ratio (*S/V*), are summarized in Table 1.

**Table 1:** Model parameters extracted from available measured data for RBCs in normal and T2DM conditions (39, 51, 55, 56, 58, 60).

|  | N-RBC | D-RBC1 | D-RBC2 | D-RBC3 |
| --- | --- | --- | --- | --- |
| $\mu$ ($\mu\text{N/m}$) | 4.73<sup>(51)</sup> | 9.46<sup>(55)</sup> | 9.46<sup>(55)</sup> | 9.46<sup>(55)</sup> |
| $S/V$ ($\mu\text{m}^2/\mu\text{m}^3$) | 1.44<sup>(60)</sup> | 1.44 | 1.04<sup>(56)</sup> | 1.04<sup>(56)</sup> |
| $\eta_m$ ($\text{pa} \cdot \text{s}$) | 0.128<sup>(39)</sup> | 0.128 | 0.128 | 0.256<sup>(58)</sup> |

The simulations were performed using a modified version of the atomistic code LAMMPS. The time integration of the motion equations is computed through a modified velocity-Verlet algorithm (35) with *λ* = 0.50 and time step Δ*t* = 1.0 *×* 10^−4^ *τ ≈* 0.66 *µs*. It takes (1.0*∼* 5.0) *×* 106 time steps for a typical simulation performed in this work.

## RESULTS AND DISCUSSION

In this section, we employ the two-component RBC model to investigate the mechanics, rheology and dynamics of T2DM RBCs. First, we probe the static and dynamic responses of T2DM RBCs subject to tensile forcing and quantify the cell deformation. Second, we simulate the dynamic behavior of T2DM RBC in shear flow and investigate the effect of membrane viscosity on the tank-treading frequency. Finally, we study the biorheological properties of individual T2DM RBCs and predict the shear viscosity of T2DM RBC suspensions.

### Mechanical properties of T2DM RBCs

To investigate the elastic response of RBCs in normal and T2DM conditions, we subject the RBC to stretching deformation by imposing an external tensile force at diametrically opposite directions which is analogous to that in optical tweezers experiments (61). In our simulations, the total stretching force, *F*_*s*_, is applied in opposite direction to ϵ *N*_*v*_ (ϵ = 0.05) vertices of the lipid bilayer component of the RBC membrane. The stretching response of the RBC is then characterized by the variation of axial (*D*_A_) and transverse (*D*_T_) diameters of the RBC (Fig. 2). For D-RBC1, we find a significant decrease in *D*_A_ when compared with that of N-RBC, which is due to the increase in shear stiffness of T2DM RBC. For D-RBC2 with a near-oblate shape, we find a further decrease in *D*_A_, indicating a further reduction in cell deformability. This result confirms that the decrease in *S/V* ratio from *∼* 1.44 to *∼* 1.04 causes a reduction in RBC deformability, which is consistent with our recent computational results that the *S/V* ratio is one of the main dominant factors in cell deformability (47). For D-RBC3, we find that the *D*_A_ values are almost exactly the same as those obtaiend for D-RBC2 (Fig. 2). This result demonstrates that the influence of the membrane viscosity on RBC stretching deformation is negligible, because it is performed under the equilibrium stretched state at a given *F*_*s*_. Hence, we conclude that the membrane shear stiffness and surface area-to-volume ratio are the two most important parameters in static RBC deformation.

**Figure 2:**
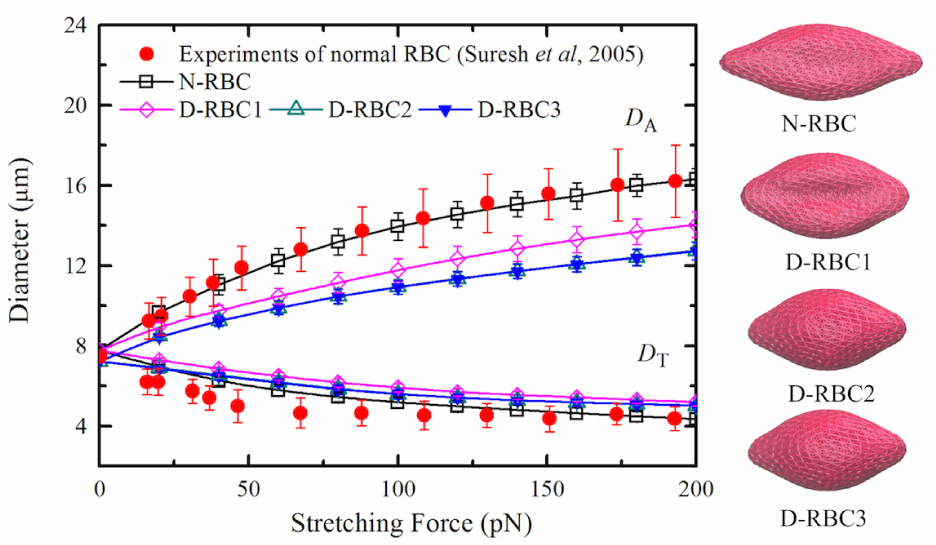
Stretching response of normal and T2DM RBC membrane at different values of the stretching force. The error bars are obtained by increasing or decreasing *µ* by 20% from their default values (Table. 1). The experimental data is adopted from (61) (Suresh *et al*, 2005), and the different stretched RBCs at stretching force *F*_*s*_ = 100 pN are presented on the right.

In addition, instantaneous membrane fluctuations of RBCs, also called “membrane flickering” (62), are commonly used to characterize the membrane stiffness. Following the work of Fedosov *et al.* (63) and Peng *et al.* (39), we then examine the membrane fluctuations of T2DM RBCs by computing instantaneous fluctuation height of the RBC membrane surface (Fig. 3). From this plot we find a good agreement between the fluctuation distributions in experiments (red line) and simulations (black square) for normal RBC. For T2DM RBCs, we find narrower distributions or smaller full-width half-maximum (FWHM) values compared to those for normal RBCs, which show the influence of local membrane curvature and effective geometry on the membrane fluctuations. These results are consistent with the previous simulations of malaria-infected RBCs at the schizont stage (63): When simulations employed a biconcave RBC shape, a wider distribution than that in experiment (62) for the schizont stage was observed; however, by employing a nearly spherical membrane (spherical RBCs at schizont stage observed in experiments) in the RBC model, the distribution became much narrower and presented a better agreement with the experiments. These computational results show that both membrane shear modulus and shape alteration play important roles in RBC membrane fluctuations. Moreover, the membrane distributions obtained from D-RBC2 and DRBC3 remain almost the same, indicating a small effect of membrane fluidity on the cell membrane fluctuations.

**Figure 3:**
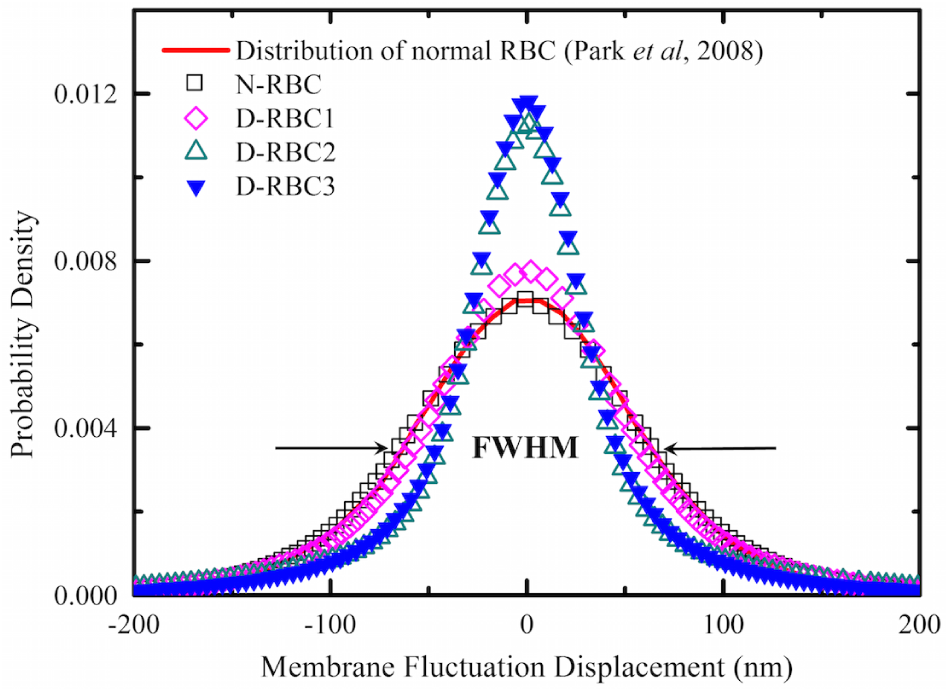
Membrane fluctuation distributions of normal and T2DM RBCs. Experimental data for membrane fluctuation distribution of normal RBC is drawn with solid red line (62) (Park *et al*, 2008). The FWHM value is about 128 nm for N-RBC, 110 nm for D-RBC1, 66 nm for D-RBC2, and 65 nm for D-RBC3.

Upon external tensile forces, a normal RBC undergoes elastic deformation, and it restores to its original state when the external force is released. Therefore, the RBC deformability can also be reflected on the shape relaxation of RBCs. It is well-accepted that the shape recovery time *t*_*c*_ ≈0.1 s for normal RBCs (51, 59, 64). When the cell age was taken into account, Williamson *et al.* found that *t*_*c*_ = 0.127 *±* 0.011 s and 0.132 *±* 0.018 s for young RBCs in normal and diabetic conditions, respectively. It increases to *t*_*c*_ = 0.152*±* 0.010 s and 0.149*±* 0.014 for old ones in healthy and diabetic cells (58). Hence, the difference in *t*_*c*_ between young and old cells is pronounced while that between normal and diabetic cells is not significant. We then perform stretching-relaxation test of normal and T2DM RBCs with different model parameters (Fig. 4). We find that the recovery processes are different from each other even though all of these modeled RBCs are able to recover their original shapes. In fact, the dynamic recovery of the RBC, *R*(*t*) = *f* (*D*_A_, *D*_T_), can be described by a time-dependent exponential decay (59, 65),

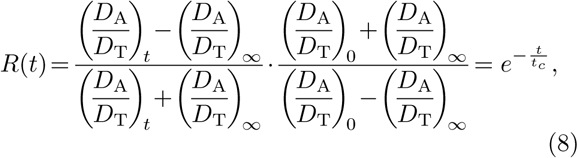

where the subscripts 0 and *∞* correspond to the ratios at *t* = 0 and *t* = *∞*. Fig. 4(b) shows the corresponding best fitting to the relaxation dynamics in Fig. 4(a). The shape recovery time is estimated to be *t*_*c*_ = 0.11, 0.08, 0.07, and 0.12 s for N-RBC, D-RBC1, D-RBC2, and D-RBC3, respectively. The comparable *t*_*c*_ value in N-RBC and D-RBC3 shows a correspondence with the experimental results (58), thus, we conclude that the effective membrane viscosity contributes largely to RBC relaxation. The underestimated *t*_*c*_ value in DRBC1 and D-RBC2 could attribute to the increased membrane shear stiffness and the reduced surface areato-volume ratio. In addition, there is no significant difference in *t*_*c*_ between D-RBC1 and D-RBC2, which is consistent with the experimental results that the geometric structure of RBC has little effect on the cell recovery process (59, 66). Our computational results on RBC mechanics from stretching behavior to recovery response demonstrate that the RBC membrane elasticity, surface area-to-volume ratio, and effective membrane viscosity are all essential to probe the RBC deformability in T2DM.

**Figure 4:**
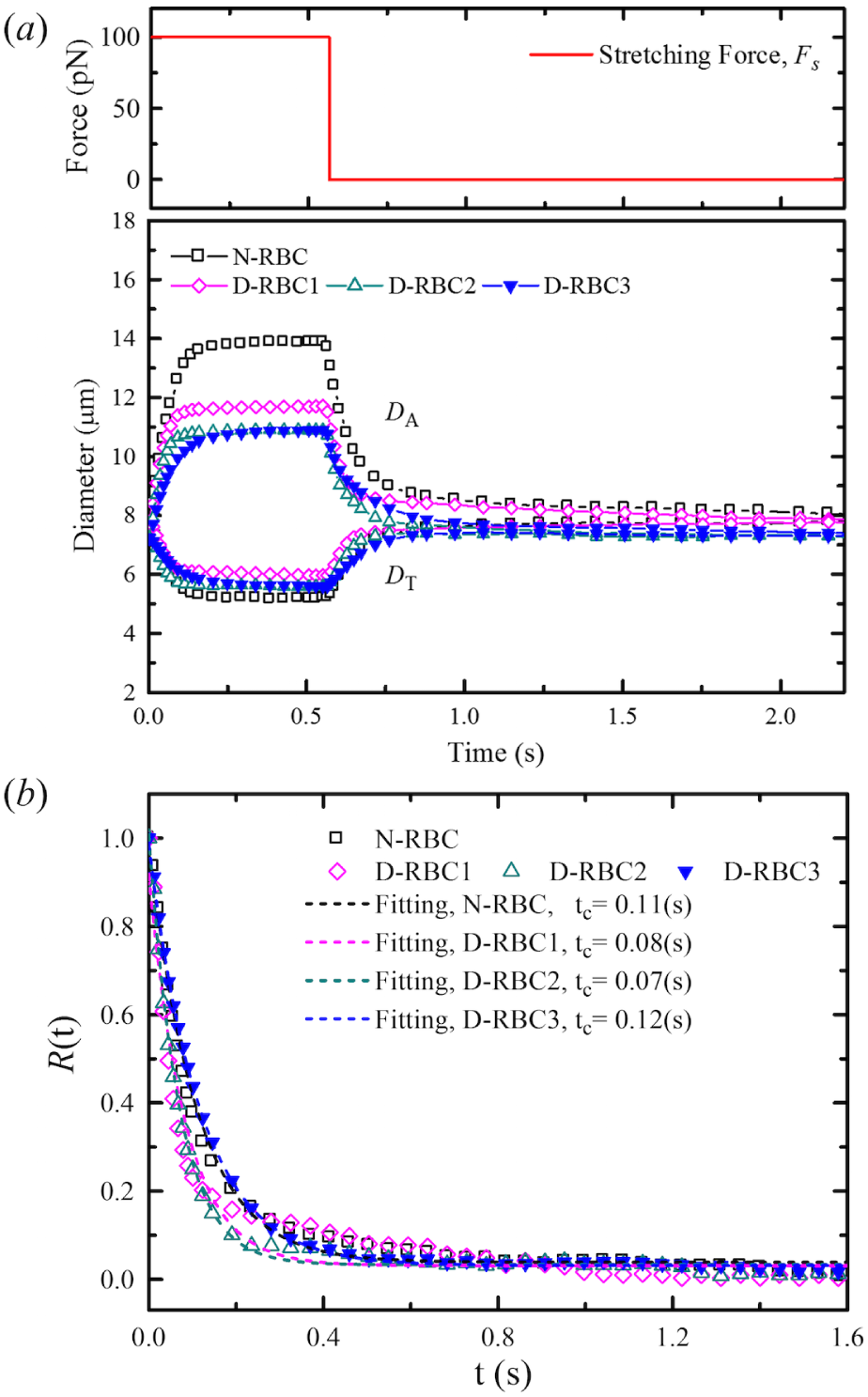
(a) Dynamic cell deformation and relaxation processes of different RBC models at the forcing shown on the top plot. (b) Estimated shape recovery time, *t*_*c*_, of different RBC models. t = 0 is the time when the external force is released and cell starts to recover its original shape.

### Dynamic behavior of individual T2DM RBC in shear flow

The RBCs in shear flow have been observed to exhibit two primary types of dynamics: tank treading (TT) motion and tumbling (TB) motion, depending on cell geometry (*S/V* ratio), cell elasticity (capillary number *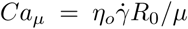* where *R*_0_ is the radius of a sphere with the same volume as an RBC), and fluid properties such as viscosity ratio (*λ* = *η*_*i*_/*η*_*o*_ where *η*_*i*_ and *η*_*o*_ are the cytoplasm and suspending fluid viscosities) (25, 67–70). At low shear rate 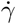, the resistance to shear causes RBCs to tumble. As 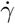 increases, the dynamics of RBCs changes from TB to TT motion (71). Physiologically, the TT motion of RBCs in flowing blood produces a lift force to prevent the RBCs from the migration toward the peripheral blood vessel, allowing high flow efficiency and sufficient oxygen transfer in blood circulation (72). In this study, we simulate the TT motion of an RBC in shear flow by placing a single RBC in linear shear flow (a Couette flow) between two planar solid walls. To produce shear flow in the fluidic channel, two solid walls move with the same speed but in opposite directions, and 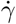 can be mediated by changing the speed of the solid walls. According to previous experimental studies (73, 74), we set *η*_*o*_ = 0.0289 *pa s* (73) and *η*_*i*_ = 0.006 *pa s* (74).

Fig. 5(a) shows the angular trajectory (*θ*) as a function of time for different RBC models at 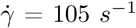 It is evident that both D-RBC1 and D-RBC2 have TT motion faster than N-RBC, which is indicated by a shorter period for the marked particle on the cell membrane to complete a TT cycle, as shown in Fig. 5(b). The accelerated TT motion for D-RBC1 than that of N-RBC is probably due to the average elongation decreases with increasing cell membrane stiffness (69). For an oblate shape of D-RBC2, it rotates even faster than DRBC1, which may owe to an increased cell thickness as reported in previous computational work of higher inclination and TT speed of osmotically swollen RBC (75). In addition, D-RBC3 exhibits a slower rotation and an extended TT period. This result shows that TT motion is sensitive to the membrane viscous dissipation, and although the swollen cell tends to speed up the TT dynamics, the elevated effective membrane viscosity slows it down even more.

**Figure 5:**
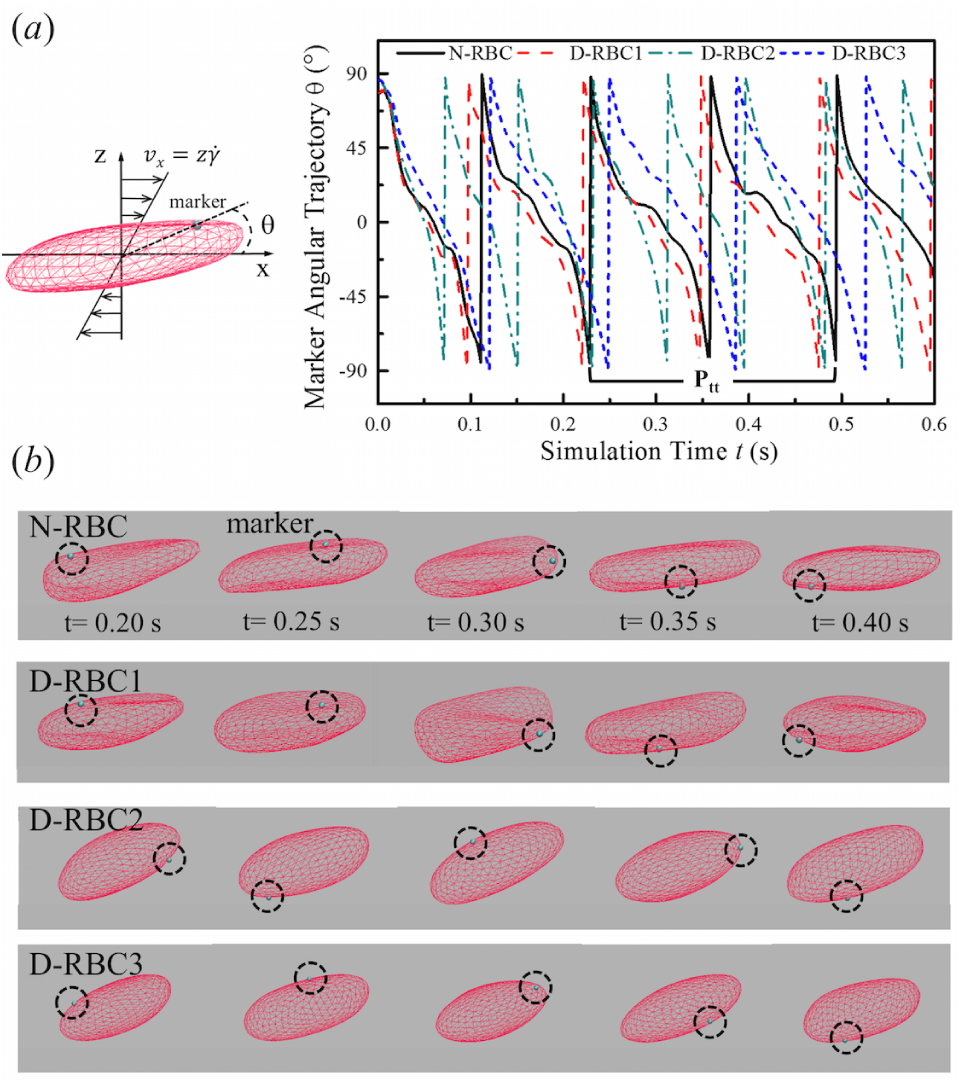
Tank treading motion of an RBC in shear flow. (a) Angular trajectory (*θ*) of a marked particle in the RBC membrane during the tank treading motion for different RBC models at shear rate 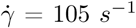. *θ* is the inclination angle between the position vector of the marked particle and flow direction. (b) Corresponding snapshots of different RBC models at *t*= 0.20, 0.25, 0.30, 0.35, and 0.40 s.

TT frequency (*f*), *i.e.* the number of TT cycles per second, is an important characteristic in the RBC TT motion. The TT frequency can be estimated from the time-dependent angular trajectories of a marked particle based on *f* = 1/*P*_*tt*_, where *P*_*tt*_ is the time for an oscillation cycle of the markered particle. In previous studies, Tran-Son-Tay *et al.* (76) and Williamson *et al.* (58) have investigated the TT frequency of RBCs in normal and diabetic conditions. They obtained a linear relationship between *f* and 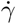 and found a slight decrease in TT frequency of diabetic RBCs compared to normal RBCs. To gain further insight into the RBC TT motion, we investigate the TT frequency of an individual RBC in normal and T2DM conditions. Fig. 6 shows that *f* increases linearly with 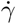 for all the four RBC models. At the same 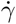, D-RBC2 has the largest *f*, N-RBC and D-RBC1 have the moderate *f*, while D-RBC3 has the smallest TT frequency. Comparing with experimental data, we find that our N-RBC and D-RBC3 have the corresponding TT frequency. These simulation results show that in addition to capturing the dynamic behavior of a normal RBC under shear flow, D-RBC3 with explicit consideration on increased effective membrane viscosity leads to an accurate model of the impaired RBC dynamics in T2DM.

**Figure 6:**
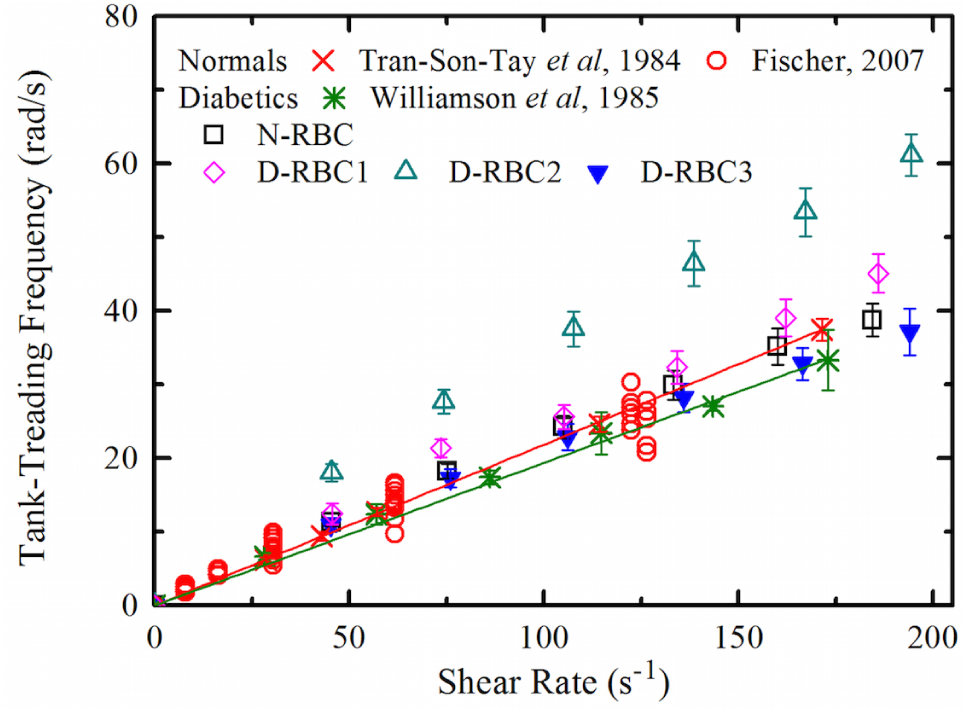
Functional dependence of RBC TT frequency with respect shear rate 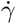. The error bars were obtained by increasing or decreasing *η*_*m*_ by 10% from their default values. Simulation results are compared with experimental data by Fischer (red circle) (73), by Tran-SonTay *et al.* (red cross) (76), and by Williamson *et al.* (green star) (58). The red line shows the linear fit line for normal RBCs in experiments, while the green line for diabetic RBCs.

### Rheological properties of T2DM RBCs in shear flow

The rheological behavior of blood samples from nondiabetic and diabetic patients have been studied in experiments (19–21, 77). Decreased cell deformability and increased cell volume in T2DM have shown significant implications for microcirculatory alterations. Computational modeling and simulations help in predicting how RBCs behave in shear flow and providing insights into how viscous flow transforms RBC shapes, and vice versa, how deformed RBCs distort the surrounding flow (24, 42, 43, 78–82). Here, in order to address the effects of cell elasticity and shape on the biorheological behavior of individual T2DM RBCs, we investigate the dynamic behavior of T2DM RBCs in a microfluidic channel with constriction. The microfluidic channel, as illustrated in the *Inlet* of Fig. 7, contains a symmetric converging and diverging nozzle shaped section. At its narrowest domain, the microchannel is 30.0 *µ*m long, 2.7 *µ*m high and has widths ranging from 4.0 to 6.0 *µ*m. The narrow and wide domains are connected by walls formed by stationary DPD particles.

**Figure 7:**
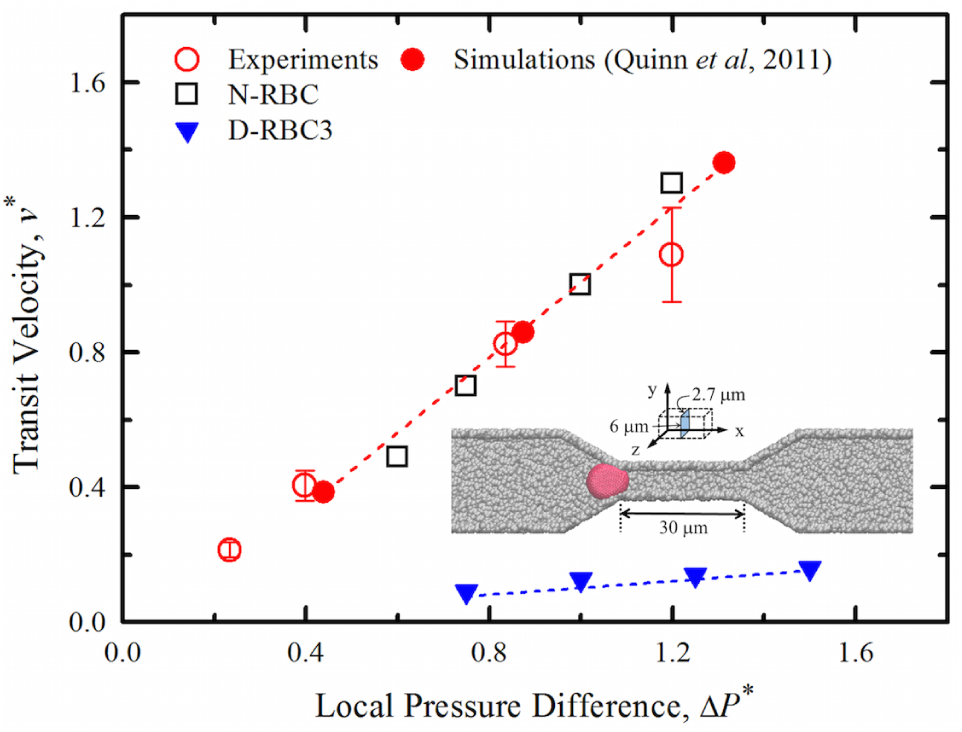
RBC traversal across a microfluidic channel with width = 6.0 *µm* at various values of pressure difference. *v*^*^ is a normalized cell transit velocity obtained through the physical velocity divided by a certain value at Δ*P* = 0.1 kPa, and Δ*P ^*^* is a normalized pressure difference obtained through the physical pressure difference divided by 0.1 kPa. Experiments (empty circle) and simulations (solid circle) of a normal RBC traversing a 6-*µm*-wide channel from Ref. (78) (Quinn *et al*, 2011) are shown. *Inset* shows the schematic of the microfluidic channel with a symmetric converging-diverging geometry.

Fig. 7 and the supporting videos S1 to S3, show the typical dynamic processes of normal and T2DM RBCs flowing through the converging-diverging microfluidic channel. Our simulations indicate that the RBCs passing through the microchannel are sensitive to *S/V* ratio (or cell volume). As shown in Fig. 7, a decrease in *S/V* ratio from 1.44 to 1.04 results in a decrease in cell transit velocity when individual RBCs travel through 6-*µm* wide microchannel. Similar to the previous computational studies of RBCs and tumor cells passing through narrow slits in Refs. (47, 83, 84), the cell volume increase (*S/V* decrease) would slow down the passing process. The average cell transit velocity decreases more significantly, however, when the individual T2DM RBCs pass through 4-*µm*-wide microchannel, see video S3. These simulations reveal that the increase of flow resistance by T2DM RBCs is larger than the resistance by normal RBCs, hence pointing to the importance of *S/V* ratio as a determinant of T2DM RBCs traversal across the narrow capillaries.

Computational models have also been proven to be an important tool in predicting the macroscopic flow properties of an RBC suspension or blood flow (e.g., yield stress and shear viscosity) from the mesoscopic properties of individual RBCs (e.g., membrane viscosity and cell deformability). Next, we employ our RBC models to examine the blood viscosity over a range of 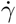 from 1 to 500 *s*^-1^. Fig. 8, and the supporting videos S4 and S5 show the relative viscosity of RBC suspensions with the N-RBC and D-RBC3 models. The model predictions clearly capture the increased blood viscosity in T2DM, in good agreement with the experimental data. This abnormality in blood viscosity is attributed to the reduced cell deformability associated with the alterations on cell shape. Our model does not include cell-cell aggregation interactions. Hence, it fails to model the formation of rouleaux structures and a tremendous viscosity increase at low shear rates (41, 85); this can be a topic for future work. Nevertheless, our model with explicit description of the RBC structure and membrane properties could lead to useful insights into the cell mechanistic processes and guide future work for better understanding the correlation between metabolic dysfunction and hematological abnormalities in T2DM.

**Figure 8:**
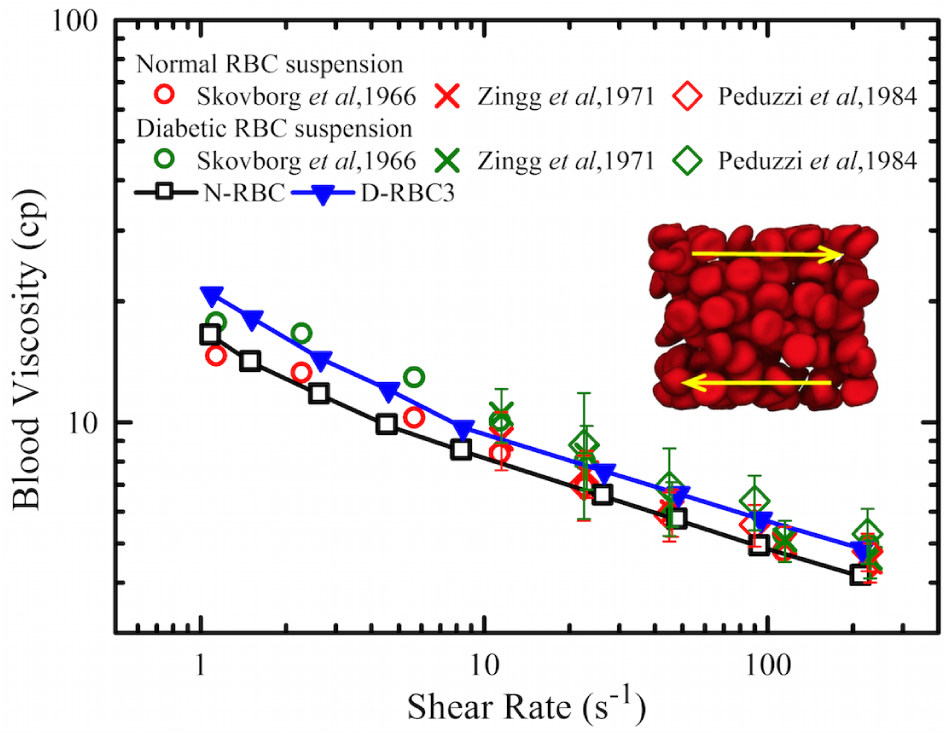
Functional dependence of shear viscosity of T2DM RBC suspension on shear rate at hematocrit *H*_*t*_ = 0.45. Experimental data are shown with circle from Ref. (19) (Skovborg *et al*, 1966), cross from Ref. (20) (Zingg *et al*, 1971), diamond from Ref. (77) (Peduzzi *et al*, 1984), and red symbols for normal RBC suspension while green symbols for diabetic RBC suspension.

## CONCLUSIONS

We have presented a two-component whole-cell model that seems to capture the biomechanical, rheological, and dynamic properties of RBCs in type 2 diabetes mellitus (T2DM) based on currently available data. The RBC membrane is modeled by two distinct networks for representing the spectrin network and the lipid bilayer. We have proposed and examined three potential T2DM RBC models (D-RBC1, D-RBC2, and DRBC3) based on the alterations in cell shape and membrane viscoelasticity measured experimentally, including increased shear stiffness, altered cell shape, and elevated membrane viscosity.

Using these RBC models, we first studied the static and dynamic deformation of T2DM RBCs subject to different tensile forcing. The significant reduction in stretching response of our diabetic RBC models against N-RBCs at the same tensile force shows the large stiffness contribution from membrane shear resistance and a near-oblate structural constraint. However, only the D-RBC3 model can exhibit reasonable relaxation processes in comparison with the corresponding experiments, indicating that membrane viscosity plays an important role in RBC deformability. Second, we quantitatively probed the linear relationship between tank-treading frequency and shear flow rate of RBCs under normal and diabetic states by our N-RBC and D-RBC3 models. The agreement of the simulation results with the currently available experiments show that the RBC complex dynamics in shear flow can be predicted by taking into account the membrane elasticity, cell geometry, and viscous dissipation. Finally, we extended the simulation from individual cell dynamics to the collective dynamics of RBC suspensions. We predicted the higher viscosity of T2DM blood than normal subjects and the results are also consistent with experimental data.

It is known that the main effect of T2DM on blood properties derives from the uncontrolled concentration of glucose. Glycated hemoglobin (HbA1c), which is formed by a simple chemical reaction between blood glucose and normal hemoglobin in RBCs, reflects the time-average blood glucose level in an individual and can be used for the diagnosis of T2DM. Experimental studies have shown that T2DM RBCs are less deformable and more fragile if there is an increased amount of HbA1c (86, 87). However, the precise effect of blood glucose level on cell deformability and blood rheology remains an open question. Our current RBC models still have limitations in facilitating the detailed exploration of diverse biophysical and biomechanical aspects involved in such cases. There is, hence, a compelling need to develop a hybrid model constructed by combining a kinetic model for the formation of HbA1c at subcelluar level with the particle-based RBC model. Such model can be used to quantify the functional relationship between glucose (or HbA1c) sensitivity and the biomechanical properties of T2DM RBCs as well as the hemorheological properties of T2DM blood. With the help of more extensive clinical tests and experimental studies, unsolved questions concerning the correlation between the degree of hyperglycemia and the alterations in diabetic hematology could be possibly addressed in the near future.

## AUTHOR CONTRIBUTIONS

Conceived and designed research: HYC XL GEK. Performed research: HYC XL. Analyzed data: HYC XL GEK. Contributed new reagents/analytic tools: HYC XL GEK. Wrote the article: HYC XL GEK.

## ACKNOWLEDGMENTS

The work described in this article was supported by the National Institutes of Health (NIH) grants U01HL114476 and U01HL116323. Part of this research was conducted using computational resources and services at the Center for Computation and Visualization (CCV), Brown University.

### CONFLICT OF INTERESTS

The authors have declared that no competing interests exist.

